# Aging Decreases the Precision of Visual Working Memory

**DOI:** 10.1101/2023.01.03.522567

**Authors:** Shahrzad Mohammadpour Esfahan, Mohammad-Hossein H.K Nili, Mehdi Sanayei, Ehsan Rezayat

**Affiliations:** Institute for Cognitive Science Studies (ICSS), Tehran, Iran; School of Electrical & Computer Engineering, College of Engineering, University of Tehran, Tehran, Iran; School of Cognitive Sciences, Institute for Research in Fundamental Sciences (IPM), Tehran, Iran; Department of Psychology and Educational Science, University of Tehran, Tehran, Iran

**Author notes:** equal contribution (Mehdi Sanayei, Ehsan Rezayat).

**Keywords:** Aging, Working Memory, Face, working memory task, Continuous error, match-to-sample

## Abstract

**Objectives:** Working memory (WM) is a cognitive ability that enables us to hold information temporarily. As age increases during life, cognitive abilities such as WM performance decrease. Errors in WM tasks arise from different sources, such as decreasing precision and random response. In the current study, we investigated the effect of age on WM and elucidated sources of errors.

**Method:** A total of 102 healthy individuals aged 18 to 71, participated in the study. A face-based visual WM task was designed and performed. Responses were collected using a graded scale in a delayed match-to-sample reproduction task.

**Results:** The error of participants increased significantly as they aged. According to our analysis of the source of error, the standard deviation of error distribution increased considerably with age. However, there was no significant change in uniform probability with age. These observations were similar between male and female participants.

**Conclusion:** We found that WM performance declines through the lifespan. Investigating the sources of error, we found that the precision of WM decreased with age. This decline was monotonous without any particular age at which a significant drop-off occurred. The results also indicated that the probability of guessing the response as a measure of random response is not affected by age.

## Introduction

Working memory (WM) refers to a mental buffer that facilitates storage. retrieval, and manipulation of information as needed for ongoing mental tasks (Hasher & Zacks, 1988). The performance of the WM deteriorates as age increases (Brockmole & Logie, 2013; Craik et al., 2010; Dobbs & Rule, 1989; Hasher & Zacks, 1988; Reinhart & Nguyen, 2019; Salthouse et al., 1991). There is a noticeable degradation of the neural substrate in the frontal lobe with age which may lead to deficits in WM in older adults (West, 1996).

Different paradigms have been used to investigate WM. In a ‘change detection’ or typical ‘delayed match-to-sample’ task where there are only correct or wrong binary answers, it is important to emphasize that just because an individual cannot recall an item, it does not mean that all information about it is lost. A correct response, on the other hand, does not reveal how well an item was retained in the memory (Zokaei & Husain, 2019). Continuing to explore the resolution with which items are retained in WM, a more recent theoretical and empirical approach has been developed (Gorgoraptis et al., 2011). Through this approach, the performance of subjects in WM is measured in a graded manner in ‘delayed-reproduction’ tasks. In this way, the precision of the response was measured, and the amount of retrieved information was analysed.

Since WM performance declines with age, it becomes important to ask whether there is an age at which WM performance declines significantly. Some studies suggested that WM abruptly declined at 50-60 years old (Brockmole & Logie, 2013; Dobbs & Rule, 1989). While other studies suggested that there is no specific age for WM decline, and as age progresses, WM gradually declines (Archer et al., 2018). Most of the studies on aging are agnostic to this issue, because they have conducted comparisons between extreme age groups (Missonnier et al., 2011; Thornton & Raz, 2006; Wild-Wall et al., 2011).

In order to clarify whether there is a specific age for WM decline, we included participants from a wide range of ages for our face WM task. We quantified the resolution with which participants recall the visual features in a graded measure. Furthermore, we investigated the effects of aging on different sources of error, the precision of the content of WM and forgetting (i.e., random guess). We found that the precision of visual memory declined with age, while age did not have any effect on random response. Our data also showed that this decline was continuous, and there was no apparent drop-off of memory precision at a certain age.

## Method

### Participants

A total of 102 healthy adults (43 females) from 18 to 71 years old participated in this experiment. Participants signed an informed consent form before the experiment. Subjects had either normal or corrected-to-normal vision. All procedures were approved by Iran University of Medical Science Ethics Committee and was in accordance with Helsinki declaration of 1964 and its revisions.

### Stimuli and Procedure

The stimuli were displayed on 15.6” Full HD laptop monitors with a 60 Hz refresh rate. The distance of subjects to the monitor was ~45 cm. A face stimulus were presented on screen and after variable delays, they were asked to reproduce the presented face (Figure 1A). We generated two faces (labeled as face A and face B in figure 1B, neutral in terms of emotion and gender, size = 5°×6°) with FaceGen Modeller 3.5 (Singular Inversions, ON, Canada). We then morphed face A to face B in 10% morph interval with InterFace software (Kramer et al., 2017). We obtained 11 morphed face stimuli in total, from 100% face A (0% face B) to 0% face A (100% face B). Morphed faces from face A to face B are shown in Figure 1B. The stimuli were displayed and subjects’ responses were collected by custom-made routines in MATLAB (MathWorks) using Psychophysics Toolbox extension 3 (Brainard, 1997; Kleiner et al., 2007; Pelli, 1997). Before the main task, an instruction was appeared on the screen and the examiner explained the procedure to the participant. After several trials of familiarization, the participant started the main task. Each trial started with a fixation spot (black, 0.3° diameter, Figure 1A). Subjects were instructed to keep fixation on it. After 1 second, a face stimulus (‘sample face’) was appeared on the screen for one second. This face stimulus was selected from 10%, 40%, 60%, or 90% morphed face A, pseudorandomly selected on each trial. After that there was a delay in which a fixation cross (white, 0.2° length of each line) was displayed at the center of the screen. Subjects were asked to keep fixation on this cross. This delay lasted for 1.5, 3, or 6 seconds, chosen pseudorandomly on each trial. After the delay, a second face (‘test face’) was presented to the subject. This ‘test face’ could by any of the morphed faces except the presented ‘sample face’ on that trial (one out of 10 possibilities). Participants were instructed to use the “Up” and “Down” arrow keys on keyboard to change the ‘test face’ through the morphs to match the ‘test face’ to the ‘sample face’. Participants pressed the “SPACE” key to submit their answers. Each block was consisted of 36 trials (3 delays × 4 sample faces × 3 repetitions). Each subject completed three blocks and they could have 1-3 minutes short break between them. We recorded subject’s choice and reaction time on each trial. We did not give participants any feedback regarding their response or reaction time.

**Figure 1.**
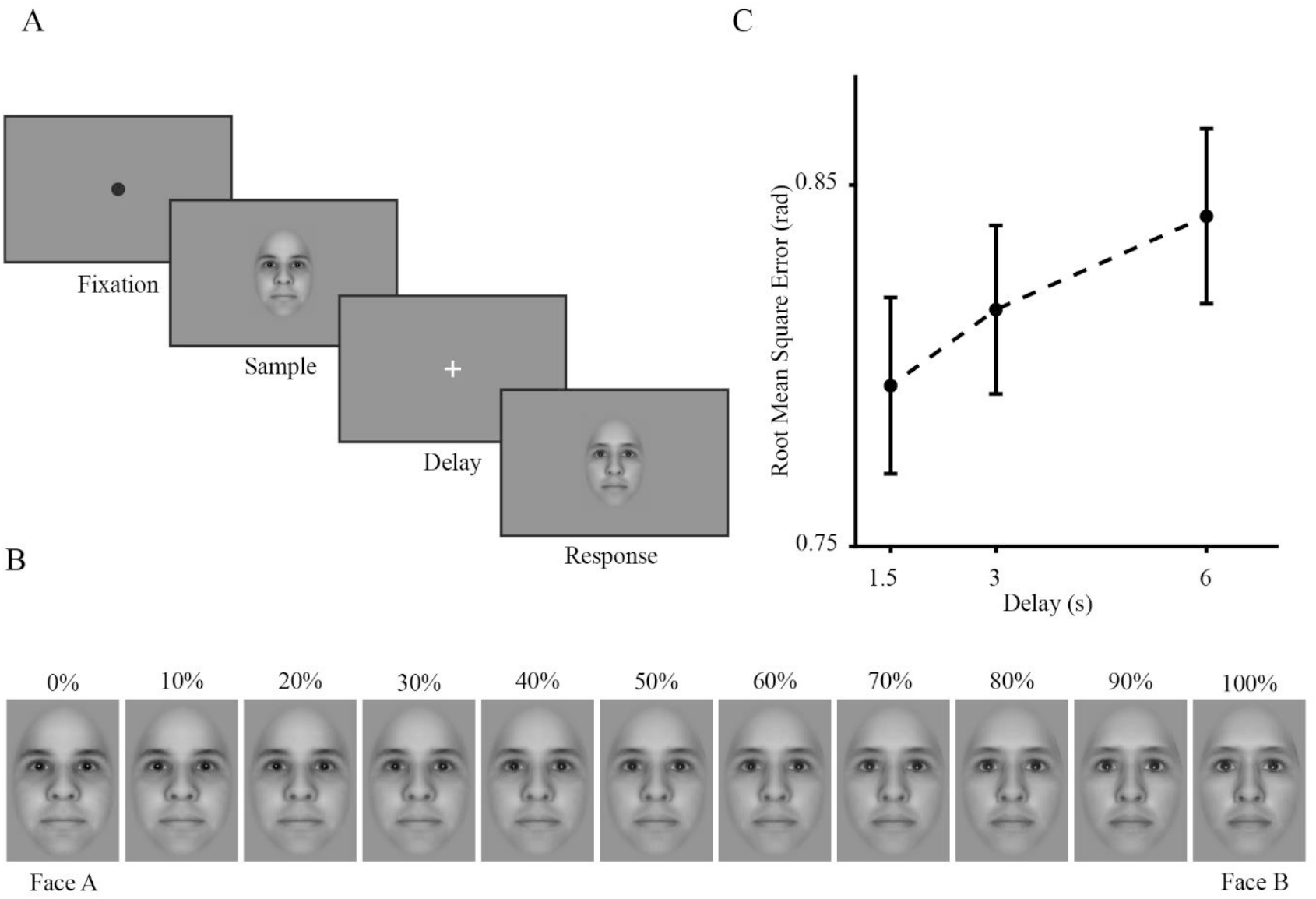
Experiment procedure and task. **(A)** A sample trial of face working memory task: Each trial began with presenting a fixation point for 1 second, followed by a ‘sample face’ presentation (1 second) and a delay (1.5, 3, or 6 seconds). A ‘test face’ was presented after the delay, and the subject had 20 seconds to change the stimulus to match it to the ‘sample face’. **(B)** Morphed faces that were used as stimuli. Faces were generated by morphing from face A to face B at 10% intervals. **(C)** The RMS error as a function of delay duration. Error bars are ± SEM (Standard Error of Mean) for each delay.

### Analysis

We defined error on each trial as the difference between the selected ‘test face’ and the presented ‘sample face’, which ranges from −10 to +10, i.e., if a subject selected a 20% morphed face A as the selected ‘test face’ while the ‘sample face’ was 60% morphed face A, the error was calculated as four (i.e., the selected ‘test face’ and the ‘sample face’ differed by four morphed images); and if the presented ‘sample face’ was 20% morphed face A and the selected ‘test face’ was 60% morphed face, the error was - 4. Due to the circular nature of possible responses, errors were coded as angular measures from – *π* to +*π* (in radians, −10 equals –*π* and +10 equal *π*).

We applied the Mixture Model developed by Bays et al. (2009) to analyse the source of error in our task. In general, on each trial, the model suggests two possible sources of error: Gaussian variability in memory for the ‘sample face’, and a fixed probability of guessing at random.

The model is described mathematically as:

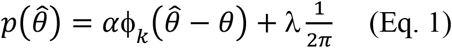

Where *θ* is the ‘sample face’ value on each trial and 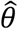 is the subject’s response (i.e., ‘test face’) in radians. α and λ are the probability of reporting the correct answer and responding randomly, respectively. *ϕ_k_* refers to the circular analogue of the Gaussian distribution (the Von Mises distribution) with mean of zero and concentration parameter κ. Maximum likelihood estimates of the parameters α, λ and κ were calculated separately for each subject and delay, using a non-linear optimization algorithm. The Von Mises κ was then converted to standard deviation. Precision was calculated as the reciprocal of the standard deviation of the error (1/SD), as in Bays and Husain (2008).

## Results

We excluded 6 participants (4 females) because they did not complete 3 blocks. The age of the included participants was 40.5 (±14) years old. As a first step, the Root Mean Square (RMS) error of each participant within each delay was calculated and a Kruskal Wallis test was conducted. Although there was not a significant difference between RMS errors as a function of delay (Kruskal Wallis test, H(2) = 2.13, p = .34), there was an upward trend as demonstrated in Figure 1C.

We then calculated the RMS error for each participant in all delays to investigate WM performance as a function of age. We found that WM impaired as age increased (r = .39, p <.001, Figure 2A). As illustrated in Figure 2B, the response time also increased significantly as age increased (r = .25, p = .02).

**Figure 2.**
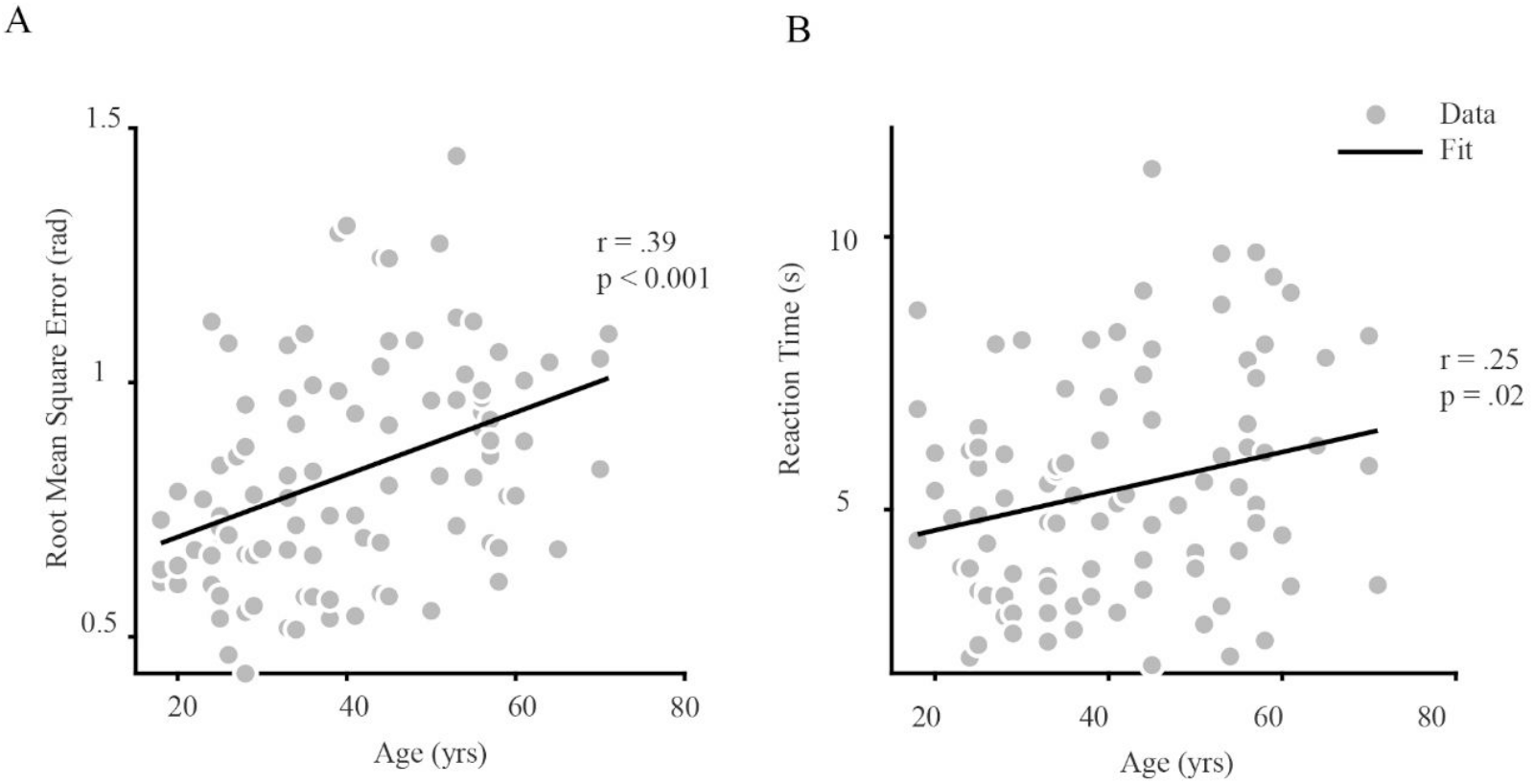
Reproduction error and reaction time increased with age. (A) The scatter plot shows the RMS error for each participant versus age. The solid line indicates the regression line. (B) The scatter plot shows the reaction time for each participant versus age. The solid line indicates the regression line. For each plot, the rho and p-value of Pearson correlation are provided.

We applied the Mixture Model developed by Bays et al. (2009) to investigate the source of error which increased as a function of age. According to the model and considering our task, there were two sources of error in our task: (1) Precision error as the standard deviation of a Von Mises distribution of recall centered on the correct answer, and (2) random guesses as uniformly distributed errors unrelated to the correct answer. As the age increased, the standard deviation of the error distribution increased significantly (r = .36, p < .001, Figure 3A). This indicates a decrease in the precision with which the ‘sample face’ was stored in the memory. However, the uniform probability of the responses was not correlated with age (*r* = .*07*, *p* = .*50*, *Figure 3B*).

**Figure 3.**
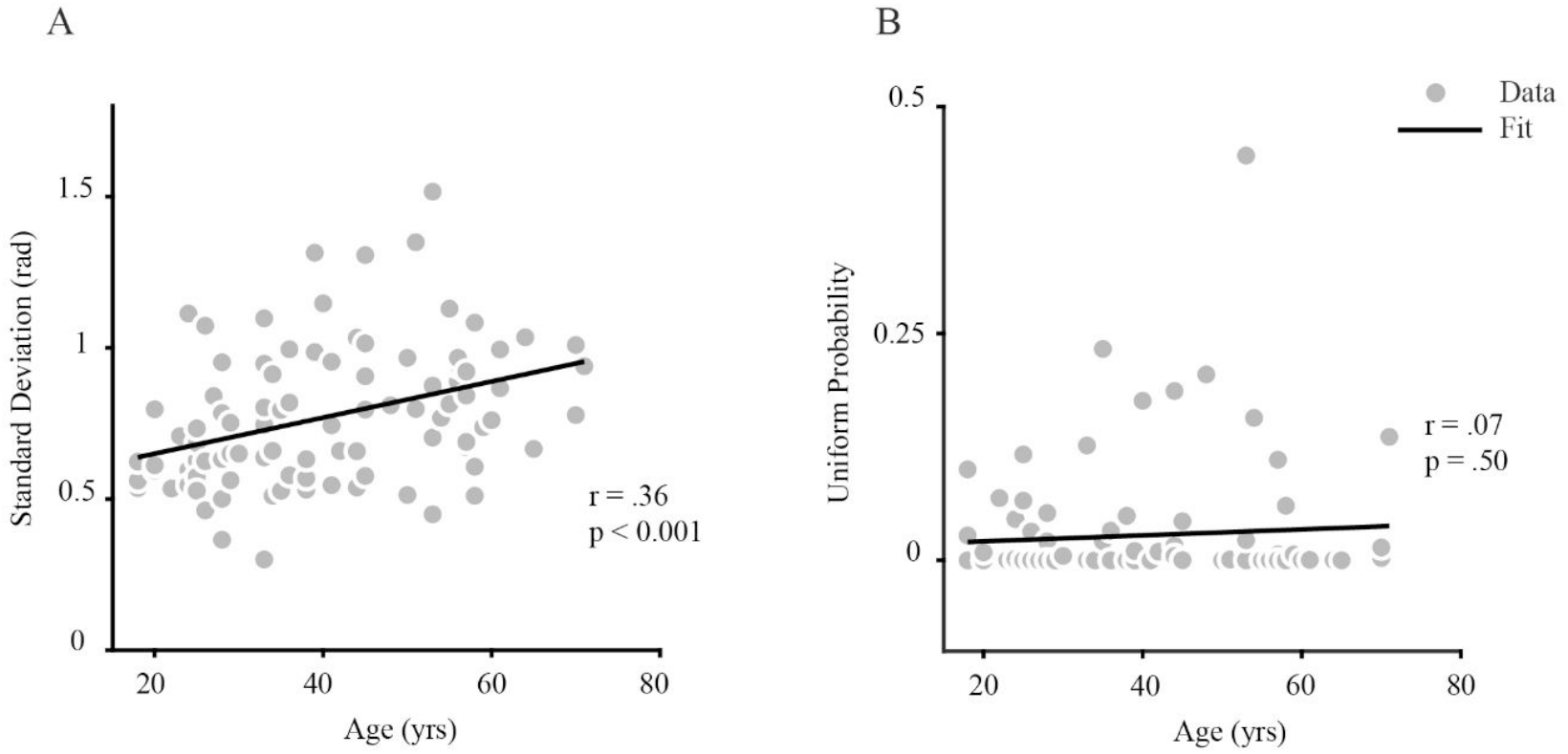
Different sources of error in the face working memory task have different relations with age. Precession error and random response error components were estimated from fitting mixture model to each participant (see Methods). **(A)** Scatter plot shows the precession error component as a function of age for each subject. Solid line shows the regression line. **(B)** Scatter plot shows the random response error component as a function of age. There was no significant relationship between the uniform probability and age. For each plot, the rho and p-value of Pearson correlation are provided.

Further investigation was conducted to determine if there was a specific age at which memory performance dropped. The participants were divided into six age groups within a 10-year bracket (and 1 year overlap between two sequential groups). Precision error and random response were tested for effects of age using a linear regression in each age group (Figure 4). The cumulative sum (CUSUM)-based approach was performed to assess the stability of the parameters at 5% level of significance. Regarding the slope of the fit to the standard deviation in each age group, the CUSUM statistics failed to reject the null hypothesis that the slope values are stable (Figure 4A). Therefore, it could be stated that there was a gradual decrease in precision as age increased without any specific age at which performance dropped. Applying the same method to the uniform probability (Figure 4B), the CUSUM statistics failed to reject stability of the slope of fitted lines with age, indicating that there was no specific age group in which the probability of random responses differed.

**Figure 4.**
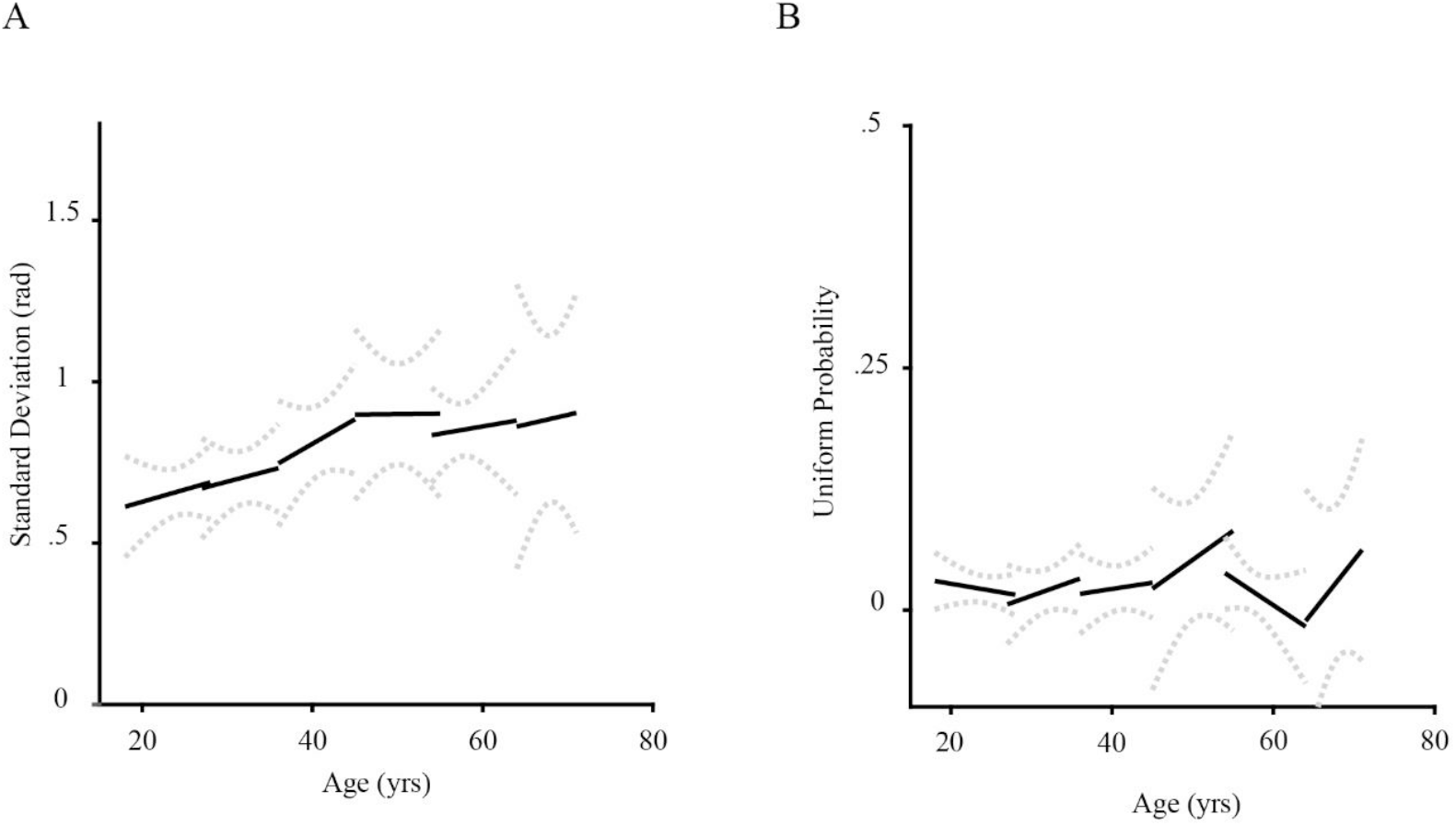
The precision of working memory decreases monotonously by age. (A) Solid lines show the fitted regression lines to the precession component of error for each age group. The dashed lines show 95% confidence interval for each age group. There is no specific age group at which precision change suddenly, indicating a monotonous rise of gaussian variability through age. (B) Solid and dashed line similar to A for random response component. The probability of random responses is not correlated with age in any of the age groups.

Next, we investigated the effect of education on our results. The mean (± sd) of education was 15.98 (± 3.54) years. We found a significant and negative correlation between the RMS error of each participant and the years of education (r = −0.40, p < .001). There was also a correlation between education years and each participant reaction time (r = −0.3, p = .006). A higher education level was associated with greater precision (r = - 0.40, p < .001). However, the probability of random response was not correlated with education (r = −0.001, p = .99). In addition, no interaction was found between age and education years (r = −0.14, p = .17).

Considering the effect of gender on WM performance through age, we found a larger effect of decline in WM in male (r = .48, p < .001) compared to the female participants (r = .35, p = .03). Both groups had a similar RMS error at age of 18 years old (intercept, .51[.33 .7], and .60[.43 .77] for male and female, respectively). RMS error increased with age with different slope (male = .008[.004 0.013], female = .004[0.001 0.008]), but still, it did not differ between males and females. The precision with which the ‘sample face’ were recalled decreased in females (r = .32, p = .04) and males (r = 0.46, p < .001), and this decline was not different between male and female (data not shown), while the probability of random response did not change significantly with age in neither (male: r = .07, p = .63; female: r = .06, p = .71).

## Discussion

In this study, we found that working memory (WM) performance declined with the increase of age in a graded WM task. We showed that precision decreased with age, while random error did not. This result is consistent with those of other studies that also indicated the decline of WM during human life span (Missonnier et al., 2011; Peich et al., 2013; Salthouse et al., 1991). In addition, we found comparable effects of gender on aging similar to previous reports (Brockmole & Logie, 2013).

Several studies have demonstrated a decline in WM performance across a wide range of age (Peich et al., 2013). They mainly used color and orientation as visual features to be recalled in WM tasks. However, in this study we generalized the well-known effect of healthy aging on WM to more complex visual stimuli (i.e., faces). In this task, we showed that the precision of the remembered face decreased as participants became older. There are two advantages using faces as to be remembered stimuli. First, given that cortical areas that code face stimuli differ from those that code color and orientation (Bae et al., 2015), our task is exploring the role of these face selective areas in WM. It has been shown that frontal areas are not the only ones that are responsible for holding information in WM. Sensory areas are actively involved in WM as well (Ester et al., 2013). So, our result suggest impairment in these face selective cortical areas as people age, as has been shown that responses of face selective areas in cortex decreases as a function of age (Zebrowitz et al., 2016). Second, a major concern in WM tasks with color and orientation is that subjects rely on canonical color categories and cardinal orientations (Pratte et al., 2017). In this study we generated two faces that were neutral in terms of both gender and emotion and morphed them. Specifically, we did not choose a photograph of a person to avoid this issue. As a result, these faces were unfamiliar we could exclude the possibility of biased answers in explaining our result.

The Mixture Model applied in this study suggests three sources of error (Bays et al., 2009). In general, these three are: gaussian variability in memory for the target, a probability of guessing at random and a certain probability of misremembering which item was the sample face. Evaluating the third source of error needs multiple items to be shown in the memory array and held in memory by subjects. In these cases, known as ‘binding errors’ or ‘swap errors’, the features of other non-probed items in the memory array can be reported instead of the features of the target. However, as we used faces as the to be remembered stimulus, using multiple faces in the memory array made the task too challenging for participants, especially elderly individuals. In that case, our choice would be to examine the WM performance of young and middle-aged participants or to simplify the task by only showing one item as the ‘sample face’ and use a life-span approach. We decided to choose the latter. As a result, despite some studies evaluating three sources of error, as the Mixture Model suggests (Peich et al., 2013), we were not able to measure swap error as a function of age. It is important to mention that Piech et al, reported that swap error indeed increased as a function of age when the memorandum was color or orientation. Whether swap error increases with age for more complex stimuli, like face, needs to be investigated.

Besides swap errors, participants can also make random errors if items are not encoded or retrieved properly on some trials, leading to guessing. This source of error that causes impairment to WM, is represented by random response. In line with a previous report, we have found that age does not affect the probability of random responses (Peich et al., 2013).

According to resource model of WM, the precision with which visual items are remembered depends on the allocation of working memory resources to them (Bays et al., 2009). Considering precision as another source of error in the Mixture Model, we showed that it decreases with age. According to our results, the deficit in WM performance is caused by less precision in older participants and not by random guesses. Considering that all participants had only one item in our memory array after omitting binding errors, and that the random response rate did not change over time, we can conclude that WM resource capacity declines as people age.

In contrast to some studies comparing two extreme age groups (Ko et al., 2014; Wild-Wall et al., 2011), this study has included participants of a wide range of ages. Therefore, the WM performance was analysed as a function of age. As a result of this approach, we were able to determine whether there was a specific age at which WM begins to decline significantly (Brockmole & Logie, 2013; Dobbs & Rule, 1989). Our study did not identify a particular age at which WM begins to deteriorate remarkably, and the decline was almost linear throughout the life span.

In addition to the mentioned results, regardless of age, all participants showed an increase in error, albeit insignificant, as the delay increased, implying the retention of visual information decays with increasing delay duration (Jonides et al., 2008). However, according to a study by Shim et al., the effect of delay duration on working memory performance was considerably weak, dependent on analysis method, and observed in only a minority of subjects.

Additionally, response times slowed as participants aged, reflecting the decrease in speed with age (Salthouse, 2000). Since the our study was unable to rule out other effects of aging such as motor delay due muscle, joint or peripheral pathology (among many other causes), we cannot claim that this effect results from a slower working memory processing speed (Birren & Fisher, 1991). More research in needed to elucidate the source of this slowing in response as people get older.

In addition, in line with previous studies, educational level positively impacted WM performance and reaction time: the higher the education, the lower the error and shorter the reaction time (Zarantonello et al., 2020). Reports from a longitudinal study suggests that education is unrelated to the rate of decline in cognitive performance. However, in a review exploring the effect of education on cognitive change in old age (Anstey & Christensen, 2000), it was reported that education has a protective effect against cognitive decline through lifespan. In future studies, longitudinal data may be carried out to determine the effect of age on WM decline as well as its impact on WM memory regardless of decline.

In line with other studies the behavioral results of the face WM task suggest less activity in fusiform face area (FFA) in older adults. However, considering the decrease of activity in prefrontal cortex (PFC) with increase of age (Fjell et al., 2010), we were not able to rule out the effect of PFC in WM performance. In future studies, activity of both PFC and FFA could be measured in face WM tasks so that the regional source of impairment could be more specified.

There were also some limitations in our study including the sample size and number of tests. Evaluating larger number of participants and conducting more face WM and cognitive tests in future studies will prepare the ground for better understanding the impairment of WM in aging. Ultimately, in future studies, it is possible to utilize brain imaging techniques, such as functional magnetic resonance imaging or measuring brain electrical activity, to investigate the WM performance in relation to the activated areas in the brain.

## Acknowledgment

We would like to thank Mahyha Moqimi and Mehran Khorasani for helping us in collecting some of the data.

## Contribution

S.M.E and E.R and M.S designed the experiment. S.M.E and M.H.K.N collected data. S.M.E and E.R analyzed the data. S.M.E and E.R and M.S prepared the manuscript.

## Data availability

The data and software code that support the findings of this study are available from corresponding authors upon reasonable request.

